# Glutamatergic and GABAergic synapses in the human spinal dorsal horn revealed with immunohistochemistry

**DOI:** 10.1101/2025.04.07.647666

**Authors:** Olivia C Davis, Andrew J Todd, Theodore J Price

## Abstract

Primary afferent neurons detect sensory stimuli in the periphery and transmit this information to the dorsal horn of the spinal cord, where it is extensively processed by excitatory and inhibitory spinal controls before being sent to higher brain centres. Immunohistochemical analysis of the synaptic architecture of these spinal circuits has been notoriously difficult in formaldehyde-fixed tissue and often requires antigen retrieval techniques to reveal antigen binding epitopes. To avoid the damage this harsh treatment can induce, studies have used antibodies raised against scaffolding proteins Homer1 and gephyrin, which anchor glutamate and GABA receptors to the membrane, respectively, to identify synaptic associations in rodent spinal cord underlying pain and itch transmission. In contrast, few studies have attempted to visualise spinal cord synapses in the human, partly due to a lack of high-quality tissue with low postmortem intervals. In this study, we reveal both excitatory and inhibitory synapses at a high resolution in human lumbar spinal cord tissue using antibodies to Homer1 and gephyrin and show that the basic organisation of these proteins within the dorsal horn is similar to that seen in the rodent. We show these postsynaptic markers are highly colocalised with glutamate and GABA receptor subunits and are in close apposition to presynaptic markers, confirming their specificity. Finally, we use Homer1-immunolabelling to demonstrate that primary afferents can form complex synaptic arrangements in human. We conclude that these antibodies can be used as reliable tools for the study of the human CNS and we have used them to reveal insight into the microanatomy of somatosensory connections in the human spinal cord that can be expanded upon in future studies.

## Introduction

Sensory stimuli in the periphery are detected by primary afferent neurons and conveyed to the dorsal horn of the spinal cord, where these primary afferents synapse onto projection neurons and interneurons and take part in complex synaptic circuits. Preclinical animal research has provided essential insight into these circuits and how the balance between excitatory and inhibitory spinal controls can influence sensory thresholds (Basbaum et al., 2009; Todd, 2010). Immunohistochemical analysis of synaptic circuits has been hindered by protein crosslinks formed during formaldehyde fixation, which can render the antigen binding epitopes of glutamate and GABA receptors inaccessible, making it difficult to identify synapses at the light microscope level (Fritschy et al., 1998; Watanabe et al., 1998). Whilst this problem can be overcome with antigen retrieval, the harsh treatment involved can affect the binding of other antibodies that may be used in combination to map out cellular networks (Gutierrez-Mecinas et al., 2016; Nagy et al., 2004).

The postsynaptic density (PSD) at both excitatory and inhibitory synapses contains receptors, together with an array of scaffolding and adapter proteins that form a dynamic signaling mesh (Sheng & Hoogenraad, 2007). Homer1 is a scaffolding protein that forms a critical part of the glutamatergic PSD and antibodies raised against Homer1 have been shown to label virtually all glutamatergic synapses in laminae I-III in the rodent spinal dorsal horn, without the need for antigen retrieval (Gutierrez-Mecinas et al., 2016). These antibodies have subsequently been used to map sensory circuits in the mouse (e.g. Albisetti et al., 2019; Boyle et al., 2019). Homer1 antibodies have also been used to identify synaptic associations in human iPSC cultures and organoids (e.g. Lutz et al., 2021; Renner et al., 2020; Teunissen et al., 2023), but they have yet to be used in studies of intact human spinal cord. Indeed, few studies have used immunohistochemistry to reveal synapses within the human spinal cord, due in part to the difficulty of accessing high quality postmortem tissue, meaning these studies are often limited by a low resolution, use tissue only from elderly patients, or are compromised by long postmortem intervals (Aousji et al., 2023; Baer et al., 2003; olde Heuvel et al., 2023).

The scaffolding protein gephyrin forms a core component of the inhibitory PSD (Tyagarajan & Fritschy, 2014) and gephyrin immunolabelling has been shown to be restricted to GABAergic, glycinergic and mixed GABA/glycinergic postsynaptic complexes in the rodent (Pfeiffer et al., 1984; Sassoè-Pognetto et al., 1995; Triller et al., 1985). Antibodies raised against the gephyrin protein have been used to study mouse inhibitory spinal circuits (Boyle et al., 2023; Davis et al., 2023) and have also been shown to produce punctate labelling that colocalises with glycine receptors in human brain and cervical spinal cord tissue (Baer et al., 2003; Waldvogel et al., 2003). It has, however, yet to be studied in association with GABA_A_ receptors or in apposition to presynaptic markers in human spinal cord tissue.

In this study, we use antibodies raised against Homer1 and gephyrin to reveal excitatory and inhibitory synapses, respectively, at a high resolution in lumbar spinal cord tissue from human organ donors ranging from 18 to 73 years of age. We also show these postsynaptic proteins are in close apposition to presynaptic markers and confirm that the basic organisation of these proteins within the dorsal horn is similar to that seen in the rodent. Finally, we use anti-Homer1 to demonstrate the complex synaptic architecture of peptide-containing primary afferents in the human dorsal horn. Together, therefore, we suggest these antibodies can be used as reliable tools to unravel the anatomy of somatosensory processing within the human spinal cord in future studies.

## Materials & Methods

### Tissue procurement

All human tissue procurement procedures were approved by the Institutional Review Board at the University of Texas at Dallas and collected in collaboration with the Southwest Transplant Alliance. Spinal cords were surgically extracted from organ donors within 1.5 – 3 hours of cross-clamp and frozen using crushed dry ice (Shiers et al., 2024). Tissue blocks were stored at −80°C until use.

### Immunohistochemistry

Lumbar spinal cord blocks from 8 organ donors were cut into 20µm transverse sections using a Leica CM1950 cryostat and mounted onto charged slides (donor demographic information can be found in **Table 1**). Multiple sections more than 60µm apart in the rostrocaudal axis were reacted for each donor. The sections were fixed in 4% freshly depolymerised formaldehyde for 15 minutes, then dehydrated for 5 minutes each in sequential concentrations of ethanol (50, 70, 100, 100%). Slides were air dried, a hydrophobic barrier drawn and the sections rehydrated in phosphate-buffered saline containing 0.3M NaCl (PBS). Sections were incubated in cocktails of primary antibodies and left overnight at 4°C in a dark, moisture-controlled tray (antibody information can be found in **Table 2**). The slides were then rinsed three times in PBS and incubated with species-specific secondary antibodies (1:500; Invitrogen), overnight at 4°C. Following a further three rinses, sections were coverslipped using Vectashield Antifade mounting medium (Vector Labs, cat# H-1000) and stored at −20°C until scanning. All antibody combinations were made in PBS containing 5% normal goat serum and 0.3% triton (PBST). For some reactions, antigen retrieval was required to visualise glutamate receptor subunits. In these cases, following fixation and dehydration, slides were submerged in pre-warmed phosphate buffer and left for 5 minutes in a shaking 37°C water bath. After this, slides were transferred to a pre-warmed solution of 0.15mg/ml pepsin (Sigma, cat #9001-7-5-6) dissolved in 0.2M hydrochloric acid and left to shake for a further 5 minutes at 37°C. Immediately after this, the slides were transferred to room temperature phosphate buffer to prevent any further pepsin activity and slides were allowed to acclimate before continuing with the antibody reaction as described above.

**Table 1:** Demographic information from donors used in this study.

| **Age (years)** | **Sex** | **Race** | **Cause of Death** | **Time post cross-clamp (hours)** |
| --- | --- | --- | --- | --- |
| 19 | Female | Hispanic | Anoxia / Drug Overdose | 1.5 |
| 45 | Female | White | Anoxia / Cardiac Arrest | 2 |
| 64 | Female | White | CVA / Stroke | 2 |
| 73 | Female | Hispanic | CVA / Stroke | 2.5 |
| 18 | Male | White | Head Trauma / GSW | 2 |
| 31 | Male | Black | Anoxia / Cardiac Arrest | 3 |
| 63 | Male | White | Head Trauma / Blunt Injury | 2.5 |
| 69 | Male | Black | CVA / Stroke | 2.5 |

**Table 2:** A list of primary antibodies used in this study.

| Antibody | Source | Catalogue # | RRID | Concentration |
| --- | --- | --- | --- | --- |
| Rabbit anti-CGRP | Immunostar | 24112 | AB_572217 | 1:500 |
| Rabbit anti-GABA<sub>A</sub><br>α2 | Synaptic<br>Systems | 224 103 | AB_2108839 | 1:250 |
| Mouse anti- gephyrin | Synaptic<br>Systems | 147 111 | AB_2232546 | 1:500 |
| Mouse anti-GluR2 | Millipore | MABN1189 | AB_2737079 | 1:250 with<br>antigen retrieval |
| Guinea pig anti-Homer1 | Synaptic<br>Systems | 160 004 | AB_10549720 | 1:500 |
| Mouse anti-NMDAR1 | Invitrogen | 32-0500 | AB_2533060 | 1:100 with<br>antigen retrieval |
| FluoTag X2 PSD-95<br>AlexaFluor 647<br>conjugate | Synaptic<br>Systems | N3702-AF647-L | AB_2936216 | 1:500 |
| Rat anti-substance P | BioRad | 8450-0505 | AB_2200292 | 1:100 |
| Rabbit anti-VGAT | Invitrogen | PA5-96231 | AB_2808033 | 1:250 |
| Rabbit anti-VGluT2 | Affinity<br>Biosciences | DF13296 | AB_2846315 | 1:250 |

For all reactions, secondary antibody control reactions were performed in parallel, following the same protocol described but with no primary antibodies added to the initial incubation to ensure there was no non-specific secondary antibody labelling. A channel, typically ∼560nM, was kept blank to scan for lipofuscin in all reactions to ensure that fluorescence in other channels was not a false positive.

### Antibody characterisation

The rabbit polyclonal CGRP antibody (ImmunoStar, #24112, used at 1:500) was raised against synthetic rat α-CGRP. Immunolabelling is prevented by pre-absorption with rat α-CGRP protein (manufacturer’s data sheet) and has been shown to produce axonal staining in human postmortem dorsal root ganglia and spinal cord tissue (Shiers et al., 2021).

The rabbit polyclonal GABA_A_ α2 antibody (Synaptic Systems, #224103, used at 1:250) was raised against a short amino acid sequence of the rat GABA_A_ receptor α2 subunit, which is also found in the human and mouse isoforms. This antibody does not produce any immunolabelling in GABA_A_ α2 subunit knock-out mice (Gao & Heldt, 2016).

The gephyrin antibody (Synaptic Systems, #147111, used at 1:500) is a mouse monoclonal antibody with an epitope corresponding to 320 amino acids found within the human, mouse and rat gephyrin protein isoforms. Punctate staining that colocalises with GABA and glycine receptor immunolabelling has been shown in human brainstem and upper cervical cord tissue (Waldvogel et al., 2003).

The GluR2 mouse monoclonal antibody (Sigma Aldrich, #MABN1189, used at 1:250) is targeted against an epitope within the N-terminal domain of the rat, mouse and human AMPA GluR2 receptor subunit isoforms. The antibody produced a single band of ∼110kDa on western blots of mouse and rat brain, as well as punctate immunoperoxidase labelling in human neocortex tissue (manufacturer’s datasheet).

The Homer1 antibody (Synaptic Systems, #160004, used at 1:500) is a polyclonal guinea pig antiserum raised against the N-terminal of the human Homer1 protein. The immunolabeling using this antibody has been shown to colocalise with other post-synaptic axon terminal markers in human brain sections (Briel et al., 2021) and iPSCs (Lutz et al., 2021).

The NMDAR1 antibody (Invitrogen, #32-0500, used at 1:100) is a monoclonal antibody raised against a fusion protein containing sequence from the intracellular loop of the NMDAR1 protein (manufacturer’s datasheet). The immunolabelling from this antibody reveals punctate staining that corresponds to the protein of NMDAR1-cDNA transfected HEK cells (Hof et al., 1996).

The FluoTag X2 PSD95 nanobody conjugated to AlexaFluor 647 (Synaptic Systems, N3702-AF647-L, used at 1:500) was raised against a region of mouse PSD95 that shares a high degree of homology with the human PSD95 isoform. This antibody has been shown to produce punctate labelling that does not overlap with other proteins of the same family with a similar structure, such as PSD93, in mouse tissue and human organoids (Kilisch et al., 2023).

The rat monoclonal substance P antibody was raised against the mammalian substance P polypeptide. The immunolabelling using this antibody can be prevented by pretreatment with synthetic substance P (Cuello et al., 1979) and has been shown to produce punctate labelling in human postmortem tissue (Mai et al., 1986).

The VGAT antibody (Invitrogen, #PA5-96231, used at 1:250) is rabbit polyclonal antibody raised against the human VGAT protein, which also shares homology with the mouse and rat isoforms. A band of ∼50kDa was shown on a western blot using mouse spinal cord (manufacturer’s datasheet).

The rabbit polyclonal VGluT2 (Affinity Biosciences, #DF13296, used at 1:250) was raised against the N-terminal domain of the human VGluT2 protein, which shares homology with the mouse and rat isoforms. A single band of ∼64kDa was shown on western blots using mouse brain and rat lung extracts (manufacturer’s datasheet).

### Confocal microscopy & Analysis

Sections were scanned with a x100 oil-immersion lens on an Olympus FV3000 confocal microscope (NA = 1.45). In all cases, the Z-interval was set to 0.3µm, the aperture was automatically set to 1 Airy unit or less, and the offset was kept at 0. All patterns were confirmed in sections from multiple donors and colocalisation analysis was conducted on Z-series captured from several sections per donor, with the data combined for each individual.

All scans were analysed using Neurolucida for Confocal software (MBF Bioscience, Williston, VT). For the analysis of glutamatergic synaptic appositions, a 5µm square grid was placed over each scan. With only the channel containing Homer1 immunolabelling visible, all Homer1-immunoreactive (-IR) puncta that contacted the gridlines were selected. The channel containing VGluT2 immunostaining was then made visible and each of these marked puncta was then assessed for the presence or absence of an apposing VGluT2-IR profile. Following this, only the VGluT2-containing channel was revealed and all respective puncta that contacted the gridlines were selected and assessed for Homer1-IR appositions. In a different reaction, Homer1-IR profiles that contacted an overlaid 5µm grid were selected and assessed for colocalisation with gephyrin-IR puncta, then in turn assessed for the presence or absence of an apposing VGAT-IR profile. In turn, all gephyrin-IR puncta that contacted gridlines were selected and assessed for colocalisation with Homer1-IR profiles, then the presence or absence of VGAT-IR appositions to assess inhibitory connections. VGAT-IR profiles were selected and assessed for the presence or absence of both gephyrin-IR and Homer1-IR postsynaptic appositions. In each case, only the channel containing the immunolabelling of interest was visible when selecting the puncta to be analysed and the other channels were only revealed after all initial selections were made.

Of the Homer1-IR and Gephyrin-IR puncta that were selected for apposition analysis, 100 were randomly chosen from multiple scans from each donor and the widest diameter in any axis measured using the Neurolucida for Confocal measure line tool.

Representative images of whole spinal cord sections were acquired using either a x10 dry lens on an Olympus FV3000RS confocal microscope or a x20 dry lens on an Olympus VS200 Slide Scanner. Some sections with Homer1 and gephyrin immunostaining were scanned using a x63 oil-immersion lens (NA = 1.4) on a Zeiss LSM 900 microscope, with a 1.4x zoom and a z-interval of 0.19µm. These Z-series were exported as a movie using Zen Blue software to show synaptic labelling in 3 dimensions (see **Supplemental Figure 1 & 2).**

### Statistical Analysis

All analysis was performed using Prism 10.0.2 Software (GraphPad, CA, USA). For individual comparisons, data from each donor was combined and then compared using either a One-way ANOVA with a Tukey multiple comparisons test, or a Kruskal-Wallis test with a Dunn’s multiple comparison post hoc test. For sex differences, the data from male and female donors were combined and compared using a Mann-Whitney test. All data are presented as mean ± SD.

## Results

An antibody raised against the Homer1 protein produces punctate immunostaining that is much more dense in the grey matter of the human spinal cord than the white matter (**Figure 1A**). Within the grey matter, there is a particularly dense band of staining in lamina II, resembling the pattern seen in rodents (Gutierrez-Mecinas et al., 2016; Figure 1A inset). These punctate Homer1-immunoreactive (Homer1-IR) profiles range from 0.25 – 2µm in length (average = 0.66 ± 0.21µm, **Figure 1B & C**), and this size does not vary significantly between individual donors of different ages and multiple causes of death (p = 0.26), or when grouped by sex (p = 0.22), suggesting the size of the PSD does not change throughout the adult lifespan. This punctate labelling appears to be consistent in size and density when viewed in consecutive optical sections scanned throughout a 20µm section (**Supplementary Figure 1**).

**Figure 1:**
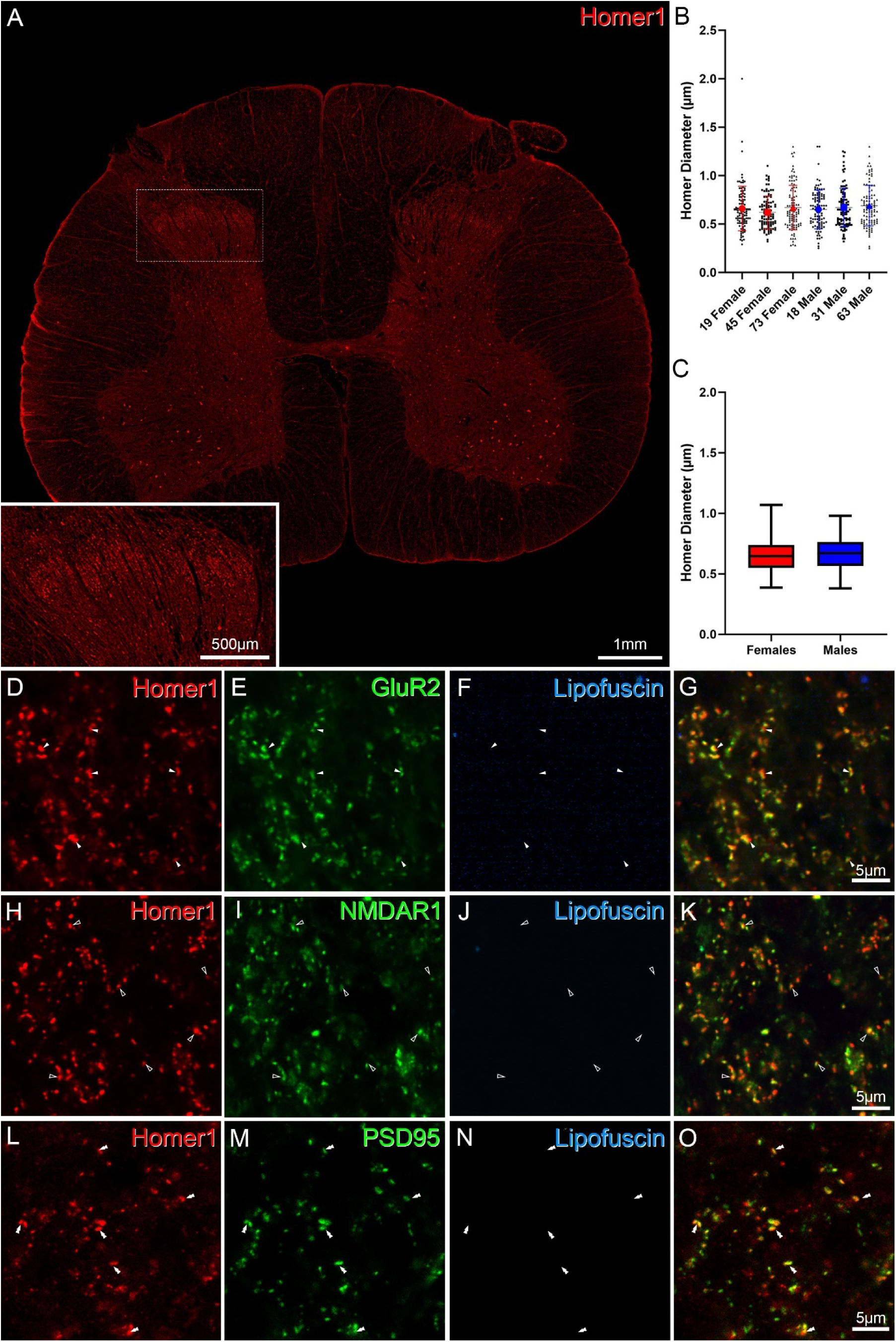
Homer1 immunolabelling in the human spinal cord is punctate and colocalises with staining for glutamate receptor subunits and PSD95. Homer1 immunolabelling forms a dense band in the superficial laminae of the human spinal cord (inset), with puncta that are 0.7µm in diameter on average throughout the grey matter (A-C). Immunolabelling for the AMPA receptor subunit GluR2 is highly colocalised with Homer1-immunoreactivity (filled arrows; D-G), as is labelling with an antibody against the NMDA NR1 receptor subunit (unfilled arrows; H-K) and labelling with a nanobody against the scaffolding protein PSD95 (double arrows; L-O).

Homer1 has been shown to crosslink several interacting proteins in the glutamatergic PSD including Shank family members (Tu et al., 1999), mGluR1 and mGluR5 (Brakeman et al., 1997; Xiao et al., 1998) and PSD95 (Tu et al., 1999). Using pepsin antigen retrieval to reveal the AMPA subunit GluR2 and the NMDA receptor subunit NMDAR1, we find that most Homer1-IR puncta are co-extensive with both of these types of ionotropic glutamate receptor subunit (**Figure 1D-K**). Similarly, there is a high degree of colocalisation between Homer1-IR profiles and immunolabelling produced using a nanobody raised against PSD95 (**Figure 1L-O**), another scaffolding protein shown to anchor ionotropic glutamate receptors to the membrane (Chen et al., 2000; Kornau et al., 1995; Tu et al., 1999). This suggests Homer1 antibodies label the post-synaptic aspect of a large proportion of glutamatergic synapses in the human spinal cord.

In the rodent, most glutamatergic presynaptic axon terminals are labelled with antibodies raised against the vesicular glutamate transporter 2 (VGluT2), as it is expressed by all excitatory spinal interneurons and most primary afferents (Häring et al., 2018; Todd et al., 2003; Zeisel et al., 2018). RNA sequencing studies suggest that this is also the case in human spinal excitatory interneurons and dorsal root ganglion neurons (Tavares-Ferreira et al., 2022; Yadav et al., 2023). Of 2637 Homer1-IR puncta selected from 6 individuals, 2029 apposed a VGluT2-IR profile (76.9 ± 2.4%, **Figure 2A-D**), with no difference between individuals (p = 0.43) or between sexes (p = 0.7). Likewise, of 2483 VGluT2-IR profiles selected, the majority apposed at least one Homer1-IR punctum (1959/ 2483; 78.9 ± 2.7%) with no individual (p = 0.29) or sex (p = 0.7) differences observed.

**Figure 2:**
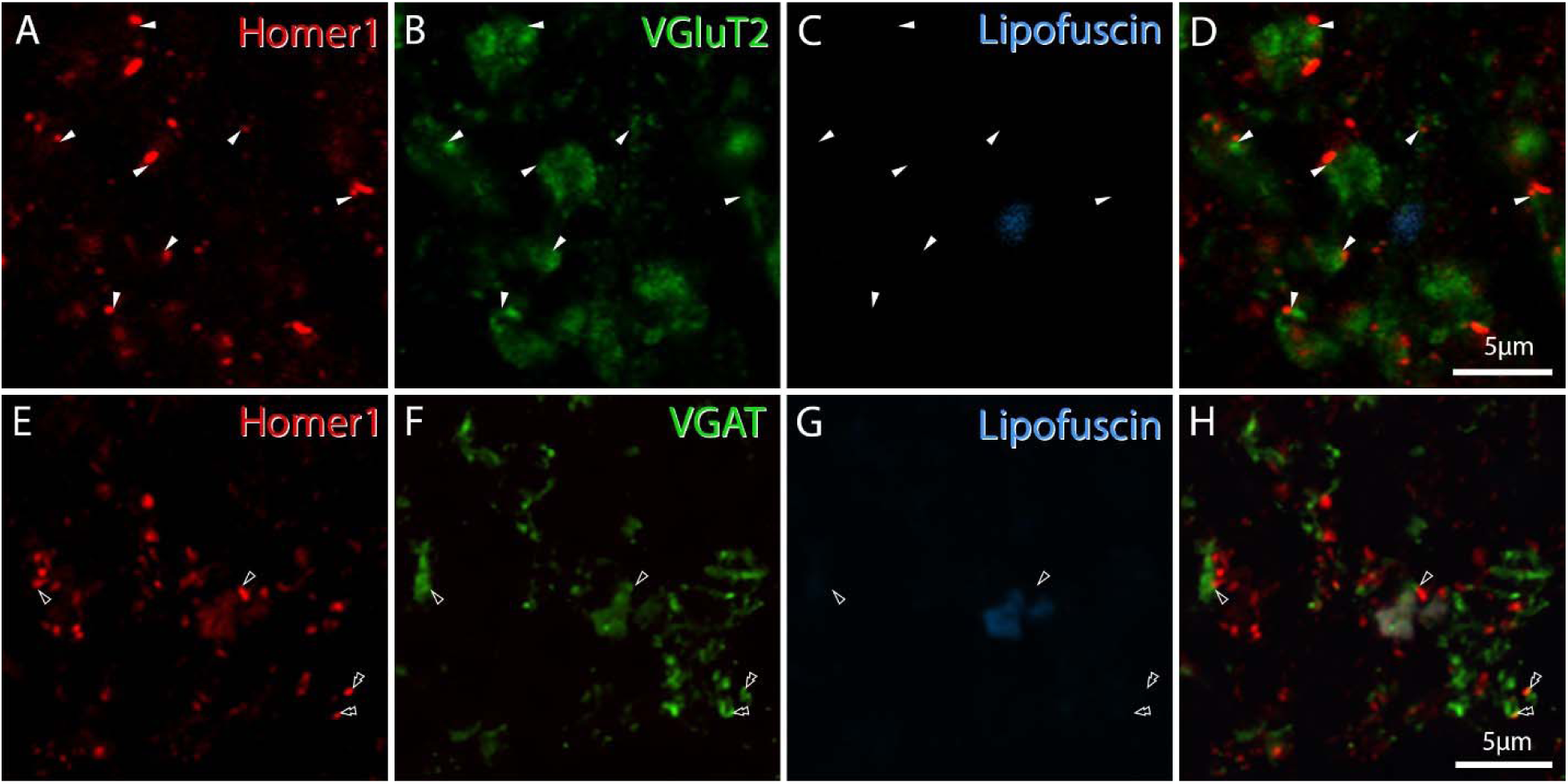
Most puncta labelled with Homer1 antibodies appose a presynaptic terminal containing VGluT2 but not VGAT. The majority of Homer1 immunolabelled profiles contact a VGluT2-immunoreactive glutamatergic terminal (filled arrows; A-D), but do not often directly appose a VGAT-immunoreactive inhibitory terminal (unfilled arrows; E-H). Occasionally Homer1 and VGAT immunoreactive profiles overlap and colocalise (double arrowhead; E-H).

In contrast, the vesicular GABA transporter (VGAT) is localised to inhibitory GABAergic and/or glycinergic presynaptic terminals. Immunolabelling for VGAT produces mostly punctate labelling in the human spinal dorsal horn (**Figure 2F)** and of 3499 Homer1-IR puncta, only 973 apposed a VGAT-IR profile (27.8 ± 3.1%, single arrow heads in **Figure 2E-H**), which was consistent between individual donors (p = 0.88) or when grouped by sex (p = 0.20). Of these Homer1/VGAT appositions, 81.3 ± 8.1% of the VGAT-IR profiles also had a gephyrin-IR apposition, suggesting these are either an excitatory and inhibitory PSD in close proximity to each other, or random contacts due to the complex synaptic arrangement of the superficial dorsal horn. Interestingly, 15.1 ± 4.1% of Homer1-IR puncta were found to either fully or partially colocalise with, rather than appose or be in contact with, VGAT-IR profiles (double arrowheads in **Figure 2E-H**) and almost 70% of these double-labelled profiles apposed a gephyrin-IR punctum (68.2 ± 1.6%; 327/486). Inhibitory spinal interneurons in rodents and non-human primates have been shown to have vesicle-containing dendrites that make dendroaxonic synapses (Carlton & Hayes, 1991; Spike & Todd, 1992). The overlap of VGAT and Homer1 immunolabelling here suggests the presence of some dendrites that are postsynaptic to glutamatergic inputs (Homer+), which also contain GABAergic and/or glycinergic synaptic vesicles, and therefore these data may reflect a similar synaptic arrangement in the human spinal cord.

Antibodies against Homer1 have been used to identify the postsynaptic targets of primary afferents terminating the dorsal horn of the rodent (e.g. Albisetti et al., 2019; Boyle et al., 2019). These antibodies have also been used to reveal the complex synaptic arrangements that some primary afferents form in lamina II, such that type I glomeruli can be seen as rings of Homer1-immunolabelled puncta (Davis et al., 2023; Gutierrez-Mecinas et al., 2016). In human, primary afferents containing calcitonin gene-related peptide often also contain substance P (Tavares-Ferreira et al., 2022) and these have been shown to terminate throughout the superficial laminae, into inner lamina II (Shiers et al., 2021). At a high magnification, the axons of primary afferents containing CGRP (with or without substance P) in lamina II were seen to be coated with Homer1-IR puncta (**Figure 3A-D**). Large swellings along these axons appeared to form complex glomerular-like synaptic structures with many Homer1-IR appositions per varicosity (asterisks and hashtags in **Figure 3A-D**), suggesting a conservation of synaptic architecture between mouse and human.

**Figure 3:**
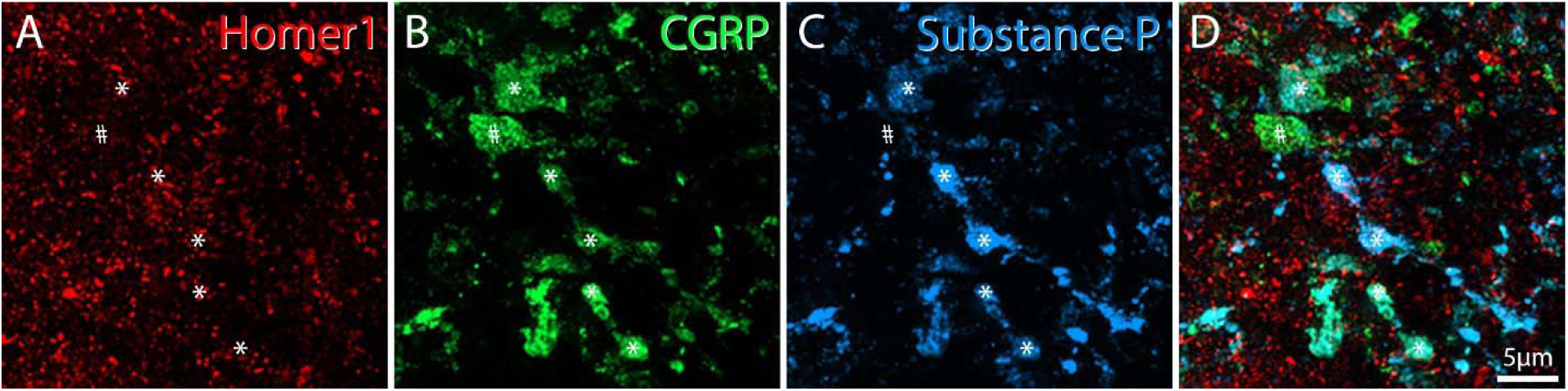
Central terminals of primary afferents containing CGRP, with or without substance P, are coated with Homer1 immunolabelled puncta. Large varicosities along peptidergic afferents appear to form glomerulus-like complex synaptic arrangements in the spinal cord. Image is a maximum intensity projection created from a short Z-series of approximately 10 optical section.

Gephyrin immunolabelling in the human spinal cord is also punctate and at a low magnification can be seen to form a dense band in lamina II (**Figure 4A and inset**). The average diameter of 600 gephyrin-IR puncta measured was 0.7 ± 0.24 µm and ranged from 0.19 – 2.1µm (**Figure 4B-C**). This was consistent between individuals (p = 0.81) and between sexes (p = 0.99). This immunostaining is also consistently bright and dense throughout consecutive optical sections taken from a 20µm tissue section (**Supplementary Figure 2**).

**Figure 4:**
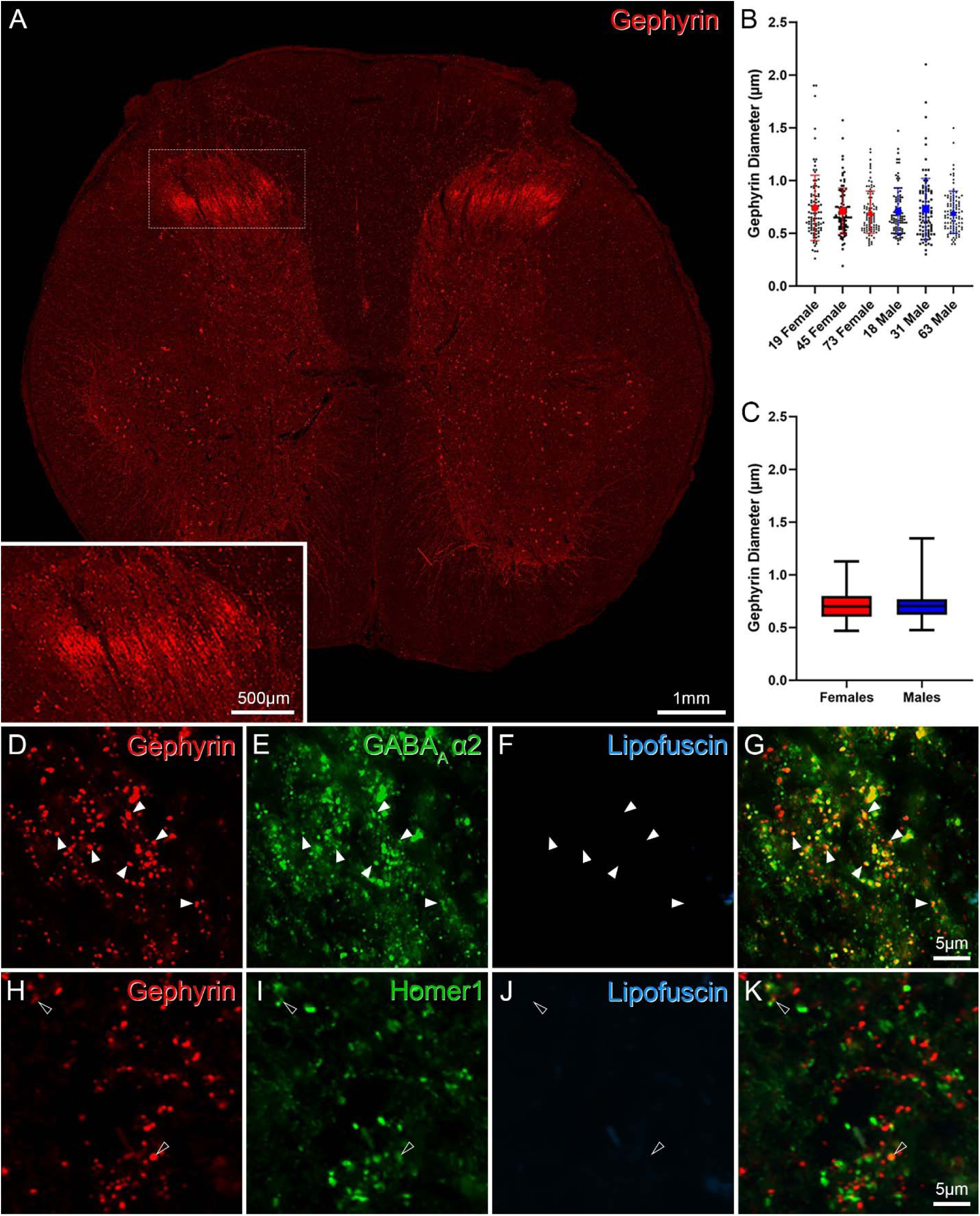
Gephyrin immunolabelling in the human spinal cord is punctate and colocalises with staining for the GABA_A_α_2_ receptor subunit, but not the glutamatergic postsynaptic protein Homer1. Gephyrin immunolabelling forms a dense band in the superficial laminae of the human spinal cord, with much less dense punctate staining elsewhere in the grey matter (A). Gephyrin-immunoreactive puncta are 0.7µm in diameter on average and do not vary in size between individuals or sex (B,C). Immunolabelling for the GABA_A_ α_2_ receptor subunit is highly colocalised with gephyrin-immunoreactivity (filled arrows; D-G) whereas Homer1-immunolabelling is rarely seen in these profiles (unfilled arrows; H-K).

Gephyrin immunolabelling has been shown to colocalise with glycine receptors in both rodent and human brain and spinal cord tissue (Baer et al., 2003; Puskár et al., 2001; Waldvogel et al., 2003), and we also find a high degree of overlap between gephyrin-IR puncta and GABA_A_-α_2_receptor subunit-IR puncta in human lumbar spinal cord (**Figure 4D – G**). Less than 10% of gephyrin-IR puncta were also immunoreactive for Homer1 (311/ 3659; 8.6 ± 4.6%; **Figure 4H - K**) and therefore gephyrin immunolabelling does not appear to label excitatory PSDs in human spinal cord tissue.

Of 3659 gephyrin-IR puncta selected from 8 individuals, 75.8 ± 4.7% apposed a VGAT-IR profile (no individual or sex difference observed, p = 0.48 and p = 0.34, respectively; **Figure 5A - D**), suggesting the majority of these appose a presynaptic inhibitory terminal. Some VGAT-IR profiles do not appose a gephyrin punctum (24.0 ± 7.2%; 762 / 3211). In the rodent, inhibitory axoaxonic synapses are formed onto the central terminals of some primary afferents, and these synapses lack gephyrin (Lorenzo et al., 2014). The VGAT-IR puncta that are not contacting a gephyrin-IR profile observed in this study may, therefore, represent axoaxonic synapses in human spinal cord.

**Figure 5:**
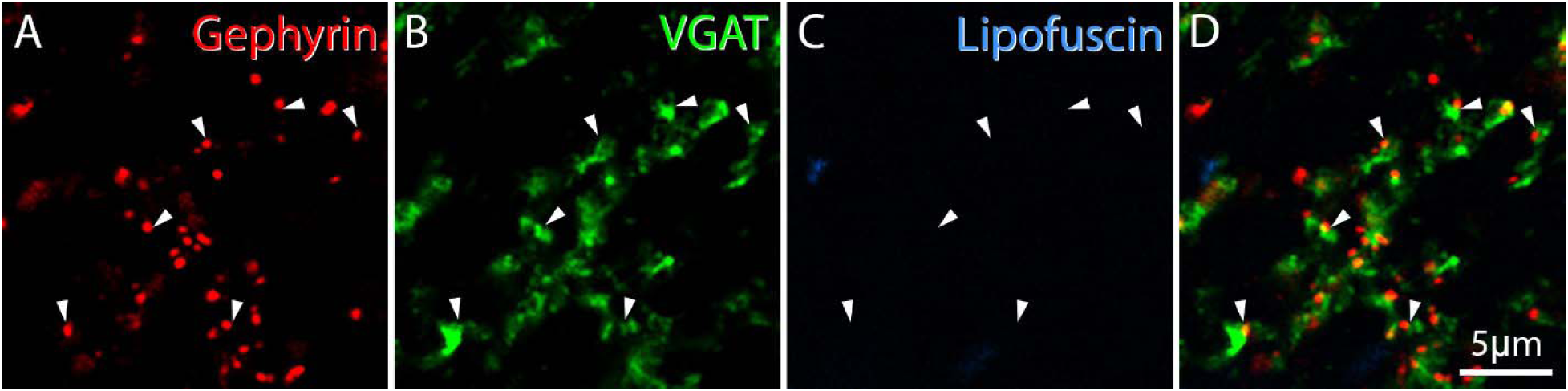
Most gephyrin immunolabelled profiles appose a VGAT immunoreactive profile. (filled arrows, A-D), suggesting the majority of gephyrin puncta are localised to inhibitory postsynaptic active zones.

## Discussion

In this study, we have shown high quality Homer1 and gephyrin immunolabelling in combination with other pre- and postsynaptic markers for the first time in intact human lumbar spinal cord sections. The general pattern of synaptic architecture appears to be consistent across species with a high density of both excitatory and inhibitory synapses in lamina II shown here in human, and in previous rodent studies (Gutierrez-Mecinas et al., 2016; Todd et al., 1995).

A high degree of colocalisation between Homer1 and both the NMDA-NR1 and AMPA GluR2 receptor subunits, as well as the anchoring protein PSD95, suggests Homer1 antibodies reliably label excitatory synapses in human spinal cord. This is also consistent with findings in the rat spinal cord. For example, Polgár et al., (2008) showed that the vast majority of excitatory synapses expressed GluR2, and Nagy et al., (2004) reported that over half of the excitatory synapses in the dorsal horn showed immunoreactivity for NMDA-NR1.

Most of the Homer1-IR puncta in human spinal cord apposed an axon terminal containing VGluT2, which is expressed by excitatory interneurons and most primary afferents terminating in the dorsal horn (Tavares-Ferreira et al., 2022; Yadav et al., 2023). Those Homer1 puncta lacking VGluT2-IR appositions may be postsynaptic to axon terminals containing VGluT1 or VGluT3, which are also expressed by a relatively small population of human primary afferents (Tavares-Ferreira et al., 2022). We observed large VGluT2-containing profiles that apposed several Homer1-IR puncta (**Figure 2A-D**), along with rings of Homer1 puncta that are a classic signature of a type I glomerulus (Davis et al., 2023) suggesting primary afferents may form complex synaptic arrangements in lamina II of the human dorsal horn. Whilst primary afferents containing CGRP project to lamina I in the rodent (Cuello et al., 1979), a previous study showed an expansion of the central projections of these primary afferents into inner lamina II in human spinal cord (Shiers et al., 2021). At a high magnification, we found that these CGRP+ axons, often also containing substance P, were coated with Homer1-IR puncta and had large varicosities that were surrounded by rings of Homer1-immunolabelled puncta. This provides further evidence that human primary afferents may form synaptic glomeruli, but a more intensive investigation is needed as only non-peptidergic C- and A∂-fibres are thought to form the central axons of glomeruli in rodents and non-human primates (Carlton et al., 1988; DiFiglia et al., 1982; Ribeiro-da-silva et al., 1989). However, because almost all human C- and A∂- sensory neurons express CGRP (Shiers et al., 2021; Tavares-Ferreira et al., 2022), this may still represent a species difference. Interestingly, many of these glomerular structures in rodent contain at least one inhibitory axoaxonic or dendroaxonic synapse (Ribeiro-Da-Silva et al., 1985), which have been shown to provide presynaptic inhibitory input to some primary afferents, including non-peptidergic C- and A-fibres (Davis et al., 2023; Hughes et al., 2012). The colocalisation of some Homer1-IR puncta with VGAT-IR profiles in this study may indicate the presence of vesicle-containing dendrites in human with the potential to form inhibitory dendroaxonic synapses as shown in the rodent, (Spike & Todd, 1992), and this could also warrant further investigation in human tissue.

Inhibitory PSDs in human postmortem spinal cord can be visualised using an antibody against the scaffolding protein gephyrin. A previous study investigating gephyrin-IR and glycine receptors in human brain and cervical spinal cord tissues observed a dense region of gephyrin staining in lamina II of the dorsal horn (Baer et al., 2003), consistent with that observed in lumbar tissue in this study. Baer et al., (2003) found many examples of dendrites in both the dorsal and ventral horn that were coated with punctate gephrin immunolabelling and estimated that approximately 60% of gephyrin-IR puncta were also immunoreactive for glycine receptors in the ventral horn of the human cervical spinal cord. Here, we observe a high degree of colocalisation between gephyrin-IR puncta and the GABA_A_-α_2_ receptor subunit in the superficial dorsal horn of lumbar spinal cord, suggesting an extensive expression of this receptor subunit at inhibitory synapses in the dorsal horn, which is consistent with the pattern found in rodent dorsal horn (Bohlhalter et al., 1996). A loss of spinal inhibition is one of the mechanisms underlying central sensitisation and abnormal sensory processing that can lead to chronic pain (Basbaum et al., 2009; Price & Prescott, 2015). Intrathecal administration of the GABA_A_ antagonist, bicuculine, has been shown to induce allodynia (Yaksh, 1989), whilst selective activation of spinal GABA_A_ receptors containing the α_2_ receptor subunit has been shown to restore sensory thresholds in inflammatory and neuropathic pain rodent models (Knabl et al., 2008). Specific positive allosteric modulators of α_2_-containing GABA_A_ channels have also been shown to effectively attenuate mechanical allodynia in both inflammatory and neuropathic rat models (Lewter et al., 2024) and may provide a novel therapeutic strategy. The data from our study suggests most GABA_A_ receptors at inhibitory PSDs in the human dorsal horn contain at least one GABA_A_-α_2_ receptor subunit and would, therefore, support the development of these targeted approaches to GABA_A_ channels containing this subunit to treat chronic pain.

In this study we have shown that synaptic labelling using these antibodies is stable and consistent between donors with various ages, sex, cause of death and postmortem intervals, meaning that these could be useful tools to start dissecting injury and disease associated changes. Using tissue from organ donors with unilateral amputations as a result of trauma or diabetic neuropathy for example, we could investigate changes in the synaptic architecture in the side of the injury compared to the contralateral side, tissue from the same donor in a region of spinal cord not associated with the affected dermatomes or tissue from donors with no history of pain. Unilateral peripheral axotomies result in the death and retraction from the spinal cord of non-peptidergic nociceptors in the rodent (Cooper et al., 2024), which is paralleled by a loss of their central terminals and output synapses in the denervated zone (Plenderleith & Snow, 1990). These antibodies, in combination with primary afferent markers, could be used to investigate whether equivalent changes or an altered distribution of either inhibitory or excitatory synapses are seen in human lumbar spinal cord following injury or disease, which could open up potential therapeutic avenues.

## Conclusion

In summary, we have shown high resolution images of both excitatory and inhibitory synaptic labelling in the human lumbar spinal dorsal horn using antibodies against pre- and postsynaptic markers. These will be valuable tools to investigate changes in synaptic architecture and circuitry in human postmortem tissue, and the approach described here should increase our understanding of human sensory processing with age and in disease states.

## Funding Statement

This research was supported by the National Institute Of Neurological Disorders And Stroke of the National Institutes of Health through the PRECISION Human Pain Network (RRID:SCR_025458), part of the NIH HEAL Initiative (https://heal.nih.gov/) under award number U19NS130608 to TJP. The content is solely the responsibility of the authors and does not necessarily represent the official views of the National Institutes of Health.

## Data availability

All data is presented within the paper. Raw image files are available upon request.

## Conflict of Interest Statement

T.J.P. is a co-founder of and holds equity in 4E Therapeutics, NuvoNuro, PARMedics, Nerveli, and Doloromics. T.J.P. has received research grants from AbbVie, Eli Lilly, Grunenthal, GSK, Evommune, Hoba Therapeutics, and The National Institutes of Health. The authors declare no conflicts of interest related to this work.

## Acknowledgements

The authors would like to thank the organ donors and their families for their invaluable gift. The authors thank the Southwest Transplant Alliance for recovery of tissues from organ donors.

## References

Albisetti, G. W., Pagani, M., Platonova, E., Hösli, L., Johannssen, H. C., Fritschy, J. M., Wildner, H., & Zeilhofer, H. U. (2019). Dorsal horn gastrin-releasing peptide expressing neurons transmit spinal itch but not pain signals. Journal of Neuroscience, 39(12), 2238–2250. 10.1523/JNEUROSCI.2559-18.2019

Aousji, O., Feldengut, S., Antonucci, S., Schön, M., Boeckers, T. M., Matschke, J., Mawrin, C., Ludolph, A. C., Del Tredici, K., Roselli, F., & Braak, H. (2023). Patterns of synaptic loss in human amyotrophic lateral sclerosis spinal cord: a clinicopathological study. Acta Neuropathologica Communications, 11(1). 10.1186/s40478-023-01616-8

Baer, K., Waldvogel, H. J., During, M. J., Snell, R. G., Faull, R. L. M., & Rees, M. I. (2003). Association of gephyrin and glycine receptors in the human brainstem and spinal cord: An immunohistochemical analysis. Neuroscience, 122(3), 773–784. 10.1016/S0306-4522(03)00543-8

Basbaum, A. I., Bautista, D. M., Scherrer, G., & Julius, D. (2009). Cellular and Molecular Mechanisms of Pain. Cell, 139(2), 267. 10.1016/J.CELL.2009.09.028

Bohlhalter, S., Weinmann, O., Mohler, H., & Fritschy, J.-M. (1996). Laminar Compartmentalization of GABA,-Receptor Subtypes in the Spinal Cord: An lmmunohistochemical Study. The Journal of Neuroscience, 16(1), 283–297.

Boyle, K. A., Gradwell, M. A., Yasaka, T., Dickie, A. C., Polgár, E., Ganley, R. P., Orr, D. P. H., Watanabe, M., Abraira, V. E., Kuehn, E. D., Zimmerman, A. L., Ginty, D. D., Callister, R. J., Graham, B. A., & Hughes, D. I. (2019). Defining a Spinal Microcircuit that Gates Myelinated Afferent Input: Implications for Tactile Allodynia. Cell Reports, 28(2), 526–540.e6. 10.1016/j.celrep.2019.06.040

Boyle, K. A., Polgár, E., Gutierrez-Mecinas, M., Dickie, A. C., Cooper, A. H., Bell, A. M., Evelline Jumolea, M., Casas-Benito, A., Watanabe, M., Hughes, D. I., Weir, G. A., Riddell, J. S., & Todd, A. J. (2023). Neuropeptide Y-expressing dorsal horn inhibitory interneurons gate spinal pain and itch signalling. ELife, 12. 10.7554/eLife.86633.2

Brakeman, P. R., Lanahan, A. A., O’Brien, R., Roche, K., Barnes, C. A., Huganir, R. L., & Worley, P. F. (1997). Homer: a protein that selectively binds metabotropic glutamate receptors. Nature, 386, 284–288. 10.1038/386284a0

Briel, N., Pratsch, K., Roeber, S., Arzberger, T., & Herms, J. (2021). Contribution of the astrocytic tau pathology to synapse loss in progressive supranuclear palsy and corticobasal degeneration. Brain Pathology, 31(4). 10.1111/bpa.12914

Carlton, S. M., & Hayes, E. S. (1991). GABAergic vesicle-containing dendrites and spines: a critical element in processing sensory input in the monkey dorsal horn. Neurosci Lett, 121, 40–42. 10.1016/0304-3940(91)90644-9

Carlton, S. M., McNeill, D. L., Chung, K., & Coggeshall, R. E. (1988). Organization of calcitonin gene-related peptide-immunoreactive terminals in the primate dorsal horn. Journal of Comparative Neurology, 276(4), 527–536. 10.1002/cne.902760407

Chen, L., Chetkovich, D. M., Petraliak, R. S., Sweeney, N. T., Kawasak^3^, Y., Wentholdk, R. J., Bredt, D. S., & Nicoll, R. A. (2000). Stargazin regulates synaptic targeting of AMPA receptors by two distinct mechanisms. Nature, 408, 936–943. www.nature.com

Cooper, A. H., Barry, A. M., Chrysostomidou, P., Lolignier, R., Wang, J., Canales, M. R., Titterton, H. F., Bennett, D. L., & Weir, G. A. (2024). Peripheral nerve injury results in a biased loss of sensory neuron subpopulations. Pain, 165(12), 2863– 2876. 10.1097/j.pain.0000000000003321

Cuello, A. C., Galfre, G., & Milstein, C. (1979). Detection of substance P in the central nervous system by a monoclonal antibody. Proceedings of the National Academy of Sciences of the United States of America, 76(7), 3532–3536. 10.1073/pnas.76.7.3532

Davis, O. C., Dickie, A. C., Mustapa, M. B., Boyle, K. A., Browne, T. J., Gradwell, M. A., Smith, K. M., Polgár, E., Bell, A. M., Kókai, É., Watanabe, M., Wildner, H., Zeilhofer, H. U., Ginty, D. D., Callister, R. J., Graham, B. A., Todd, A. J., & Hughes, D. I. (2023). Calretinin-expressing islet cells are a source of pre- and post-synaptic inhibition of non-peptidergic nociceptor input to the mouse spinal cord. Scientific Reports, 13(1). 10.1038/s41598-023-38605-9

DiFiglia, M., Aronin, N., & Leeman, S. E. (1982). Light microscopic and ultrastructural localization of immunoreactive substance P in the dorsal horn of monkey spinal cord. Neuroscience, 7(5), 1127–1139. 10.1016/0306-4522(82)91120-4.

Fritschy, J. M., Weinmann, O., Wenzel, A., & Benke, D. (1998). Synapse-specific localization of NMDA and GARA(A) receptor subunits revealed by antigen-retrieval immunohistochemistry. Journal of Comparative Neurology, 390(2), 194–210. 10.1002/(SICI)1096-9861(19980112)390:2<194::AID-CNE3>3.0.CO;2-X

Gao, Y., & Heldt, S. A. (2016). Enrichment of GABAA receptor α-subunits on the axonal initial segment shows regional differences. Frontiers in Cellular Neuroscience, 10. 10.3389/fncel.2016.00039

Gutierrez-Mecinas, M., Kuehn, E. D., Abraira, V. E., Polgár, E., Watanabe, M., & Todd, A. J. (2016). Immunostaining for Homer reveals the majority of excitatory synapses in laminae I-III of the mouse spinal dorsal horn. Neuroscience, 329, 171–181. 10.1016/j.neuroscience.2016.05.009

Häring, M., Zeisel, A., Hochgerner, H., Rinwa, P., Jakobsson, J. E. T., Lönnerberg, P., La Manno, G., Sharma, N., Borgius, L., Kiehn, O., Lagerström, M. C., Linnarsson, S., & Ernfors, P. (2018). Neuronal atlas of the dorsal horn defines its architecture and links sensory input to transcriptional cell types. Nature Neuroscience, 21(6), 869–880. 10.1038/s41593-018-0141-1

Hof, P. R., Vissavajjhala, P., Rosenthal, R. E., Fiskum, G., & Morrison, J. H. (1996). Distribution of glutamate receptor subunit proteins GluR2(4), GluR5/6/7, and NMDAR1 in the canine and primate cerebral cortex: a comparative immunohistochemical analysis. Brain Research, 723, 77–89.

Hughes, D. I., Sikander, S., Kinnon, C. M., Boyle, K. A., Watanabe, M., Callister, R. J., & Graham, B. A. (2012). Morphological, neurochemical and electrophysiological features of parvalbumin-expressing cells: A likely source of axo-axonic inputs in the mouse spinal dorsal horn. Journal of Physiology, 590(16), 3927–3951. 10.1113/jphysiol.2012.235655

Kilisch, M., Gere-Becker, M., Wüstefeld, L., Bonnas, C., Crauel, A., Mechmershausen, M., Martens, H., Götzke, H., Opazo, F., & Frey, S. (2023). Simple and Highly Efficient Detection of PSD95 Using a Nanobody and Its Recombinant Heavy-Chain Antibody Derivatives. International Journal of Molecular Sciences, 24(8). 10.3390/ijms24087294

Knabl, J., Witschi, R., Hösl, K., Reinold, H., Zeilhofer, U. B., Ahmadi, S., Brockhaus, J., Sergejeva, M., Hess, A., Brune, K., Fritschy, J. M., Rudolph, U., Möhler, H., & Zeilhofer, H. U. (2008). Reversal of pathological pain through specific spinal GABAA receptor subtypes. Nature, 451(7176), 330–334. 10.1038/nature06493

Kornau, H.-C., Schenker, L. T., Kennedy, M. B., & Seeburg, P. H. (1995). Domain Interaction Between NMDA Receptor Subunits and the Postsynaptic Density Protein PSD-95. Science, 269(5231), 1737–1740. https://www.science.org

Lewter, L. A., Woodhouse, K., Tiruveedhula, V. V. N. P. B., Cook, J. M., & Li, J.-X. (2024). Antinociceptive effects of α2/α3-subtype selective GABA A receptor positive allosteric modulators KRM-II-81 and NS16085 in rats: behavioral specificity . *Journal of Pharmacology and Experimental Therapeutics*, JPET-AR-2023–002070. 10.1124/jpet.123.002070

Lorenzo, L. E., Godin, A. G., Wang, F., St-Louis, M., Carbonetto, S., Wiseman, P. W., Ribeiro-da-Silva, A., & De Koninck, Y. (2014). Gephyrin clusters are absent from small diameter primary afferent terminals despite the presence of GABAA receptors. Journal of Neuroscience, 34(24), 8300–8317. 10.1523/JNEUROSCI.0159-14.2014

Lutz, A. K., Pérez Arévalo, A., Ioannidis, V., Stirmlinger, N., Demestre, M., Delorme, R., Bourgeron, T., & Boeckers, T. M. (2021). SHANK2 Mutations Result in Dysregulation of the ERK1/2 Pathway in Human Induced Pluripotent Stem Cells-Derived Neurons and Shank2(−/−) Mice. Frontiers in Molecular Neuroscience, 14. 10.3389/fnmol.2021.773571

Mai, J. K., Stephens, $ P H, Hopf, ? A, & Cuellot, A. C. (1986). Substance P in the Human Brain. Neuroscience, 17(3), 709–739.

Nagy, G. G., Al-Ayyan, M., Andrew, D., Fukaya, M., Watanabe, M., & Todd, A. J. (2004). Widespread expression of the AMPA receptor GluR2 subunit at glutamatergic synapses in the rat spinal cord and phosphorylation of GluR1 in response to noxious stimulation revealed with an antigen-unmasking method. Journal of Neuroscience, 24(25), 5766–5777. 10.1523/JNEUROSCI.1237-04.2004

Nagy, G., Watanabe, M., Fukaya, M., & Todd, A. J. (2004). Synaptic distribution of the NR1, NR2A and NR2B subunits of the N-methyl-D-aspartate receptor in the rat lumbar spinal cord revealed with an antigen-unmasking technique. European Journal of Neuroscience, 20(12), 3301–3312. 10.1111/j.1460-9568.2004.03798.x

olde Heuvel, F., Ouali Alami, N., Aousji, O., Pogatzki-Zahn, E., Zahn, P. K., Wilhelm, H., Deshpande, D., Khatamsaz, E., Catanese, A., Woelfle, S., Schön, M., Jain, S., Grabrucker, S., Ludolph, A. C., Verpelli, C., Michaelis, J., Boeckers, T. M., & Roselli, F. (2023). Shank2 identifies a subset of glycinergic neurons involved in altered nociception in an autism model. Molecular Autism, 14(1). 10.1186/s13229-023-00552-7

Pfeiffer, F., Simler, R., Grenningloh, G., & Betz, H. (1984). Monoclonal antibodies and peptide mapping reveal structural similarities between the subunits of the glycine receptor of rat spinal cord. Proceedings of the National Academy of Sciences, 81(22), 7224–7227. 10.1073/pnas.81.22.7224

Plenderleith, M. B., & Snow, P. J. (1990). The effect of peripheral nerve section on lectin binding in the superficial dorsal horn of the rat spinal cord. Brain Research, 507(1), 146–150. 10.1016/0006-8993(90)90534-I

Polgár, E., Watanabe, M., Hartmann, B., Grant, S. G. N., & Todd, A. J. (2008). Expression of AMPA receptor subunits at synapses in laminae I-III of the rodent spinal dorsal horn. Molecular Pain, 4. 10.1186/1744-8069-4-5

Puskár, Z., Polgár, E., & Todd, A. J. (2001). A population of large lamina I projection neurons with selective inhibitory input in rat spinal cord. Neuroscience, 102(1), 167–176. 10.1016/S0306-4522(00)00445-0

Renner, H., Grabos, M., Becker, K. J., Kagermeier, T. E., Wu, J., Otto, M., Peischard, S., Zeuschner, D., Tsytsyura, Y., Disse, P., Klingauf, J., Leidel, S. A., Seebohm, G., Schöler, H. R., & Bruder, J. M. (2020). A fully automated high-throughput workflow for 3d-based chemical screening in human midbrain organoids. ELife, 9, 1–39. 10.7554/eLife.52904

Ribeiro-Da-Silva, A., Pignatelli, D., & Coimbra, A. (1985). Synaptic architecture of glomeruli in superficial dorsal horn of rat spinal cord, as shown in serial reconstructions. Journal of Neurocytology, 14(2), 203–220. 10.1007/BF01258448

Ribeiro-da-silva, A., Tagari, P., & Cuello, A. C. (1989). Morphological characterization of substance P-like immunoreactive glomeruli in the superficial dorsal horn of the rat spinal cord and trigeminal subnucleus caudalis: A quantitative study. Journal of Comparative Neurology, 281(4), 497–515. 10.1002/cne.902810402

Sassoè-Pognetto, M., Kirsch, J., Grünert, U., Greferath, U., Fritschy, J. M., Möhler, H., Betz, H., & Wässle, H. (1995). Colocalization of gephyrin and GABAA-receptor subunits in the rat retina. Journal of Comparative Neurology, 357(1), 1–14. 10.1002/cne.903570102

Sheng, M., & Hoogenraad, C. C. (2007). The postsynaptic architecture of excitatory synapses: A more quantitative view. Annual Review of Biochemistry, 76, 823–847. 10.1146/annurev.biochem.76.060805.160029

Shiers, S., Saad Yousuf, M., Mwirigi, J., Cervantes, A., & Price, T. (2024). Human Ganglia and Spinal Cord Tissue Procurement from Organ Donors and Tissue Quality Assessment v1. In protocols.io. protocols.io. 10.17504/protocols.io.kqdg32qr1v25/v1

Shiers, S., Sankaranarayanan, I., Jeevakumar, V., Cervantes, A., Reese, J. C., & Price, T. J. (2021). Convergence of peptidergic and non-peptidergic protein markers in the human dorsal root ganglion and spinal dorsal horn. Journal of Comparative Neurology, cne.25122. 10.1002/cne.25122

Spike, R. C., & Todd, A. J. (1992). Ultrastructural and immunocytochemical study of lamina II islet cells in rat spinal dorsal horn. Journal of Comparative Neurology, 323(3), 359–369. 10.1002/cne.903230305

Tavares-Ferreira, D., Shiers, S., Ray, P. R., Wangzhou, A., Jeevakumar, V., Sankaranarayanan, I., Cervantes, A. M., Reese, J. C., Chamessian, A., Copits, B. A., Dougherty, P. M., Gereau IV, R. W., Burton, M. D., Dussor, G., & Price, T. J. (2022). Spatial transcriptomics of dorsal root ganglia identifies molecular signatures of human nociceptors. Science Translational Medicine, 14(632). 10.1126/scitranslmed.abj8186

Teunissen, M. W. A., Lewerissa, E., Van Hugte, E. J. H., Wang, S., Ockeloen, C. W., Koolen, D. A., Pfundt, R., Marcelis, C. L. M., Brilstra, E., Howe, J. L., Scherer, S. W., Le Guillou, X., Bilan, F., Primiano, M., Roohi, J., Piton, A., De Saint Martin, A., Baer, S., Seiffert, S., … Nadif Kasri, N. (2023). ANK2 loss-of-function variants are associated with epilepsy, and lead to impaired axon initial segment plasticity and hyperactive network activity in hiPSC-derived neuronal networks. Human Molecular Genetics, 32(14), 2373–2385. 10.1093/hmg/ddad081

Todd, A. J. (2010). Neuronal circuitry for pain processing in the dorsal horn. Nature Reviews Neuroscience, 11(12), 823–836. 10.1038/nrn2947

Todd, A. J., Hughes, D. I., Polgár, E., Nagy, G. G., Mackie, M., Ottersen, O. P., & Maxwell, D. J. (2003). The expression of vesicular glutamate transporters VGLUT1 and VGLUT2 in neurochemically defined axonal populations in the rat spinal cord with emphasis on the dorsal horn. European Journal of Neuroscience, 17(1), 13–27. 10.1046/j.1460-9568.2003.02406.x

Todd, A. J., Spike, R. C., Chong, D., & Neilson, M. (1995). The Relationship Between Glycine and Gephyrin in Synapses of the Rat Spinal Cord. European Journal of Neuroscience, 7(1), 1–11. 10.1111/j.1460-9568.1995.tb01014.x

Triller, A., Cluzeaud, O., Pfeiffer, F., Betz, H., & Korn, H. (1985). Distribution of Glycine Receptors at Central Synapses: An Immunoelectron Microscopy Study. The Journal of Cell Biology, 101, 683–688. 10.1083/jcb.101.2.683

Tu, J. C., Xiao, B., Naisbitt, S., Yuan, J. P., Petralia, R. S., Brakeman, P., Doan, A., Aakalu, V. K., Lanahan, A. A., Sheng, M., & Worley, P. F. (1999). Coupling of mGluR/Homer and PSD-95 Complexes by the Shank Family of Postsynaptic Density Proteins. Neuron, 23, 583–592. 10.1016/s0896-6273(00)80810-7

Tyagarajan, S. K., & Fritschy, J. M. (2014). Gephyrin: A master regulator of neuronal function? Nature Reviews Neuroscience, 15(3), 141–156. 10.1038/nrn3670

Waldvogel, H. J., Baer, K., Snell, R. G., During, M. J., Faull, R. L. M., & Rees, M. I. (2003). Distribution of gephyrin in the human brain: an immunohistochemical analysis. Neuroscience, 116(1), 145–156. 10.1016/s0306-4522(02)00550-x

Watanabe, M., Fukaya, M., Sakimura, K., Manabe, T., Mishina, M., & Inoue, Y. (1998). Selective scarcity of NMDA receptor channel subunits in the stratum lucidum (mossy fibre-recipient layer) of the mouse hippocampal CA3 subfield. European Journal of Neuroscience, 10(2), 478–487. 10.1046/j.1460-9568.1998.00063.x

Xiao, B., Tu, J. C., Petralia, R. S., Yuan, J. P., Doan, A., Breder, C. D., Ruggiero, A., Lanahan, A. A., Wenthold, R. J., & Worley, P. F. (1998). Homer Regulates the Association of Group 1 Metabotropic Glutamate Receptors with Multivalent Complexes of Homer-Related, Synaptic Proteins. Neuron, 21(4), 707–716. 10.1016/s0896-6273(00)80588-7

Yadav, A., Matson, K. J. E., Li, L., Hua, I., Petrescu, J., Kang, K., Alkaslasi, M. R., Lee, D. I., Hasan, S., Galuta, A., Dedek, A., Ameri, S., Parnell, J., Alshardan, M. M., Qumqumji, F. A., Alhamad, S. M., Wang, A. P., Poulen, G., Lonjon, N., … Levine, A. J. (2023). A cellular taxonomy of the adult human spinal cord. Neuron, 111(3), 328–344.e7. 10.1016/j.neuron.2023.01.007

Yaksh, T. L. (1989). Behavioral and autonomic correlates of the tactile evoked allodynia produced by spinal glycine inhibition: effects of modulatory receptor systems and excitatory amino acid antagonists. Pain, 37(1), 111–123. 10.1016/0304-3959(89)90160-7

Zeisel, A., Hochgerner, H., Lönnerberg, P., Johnsson, A., Memic, F., van der Zwan, J., Häring, M., Braun, E., Borm, L. E., La Manno, G., Codeluppi, S., Furlan, A., Lee, K., Skene, N., Harris, K. D., Hjerling-Leffler, J., Arenas, E., Ernfors, P., Marklund, U., & Linnarsson, S. (2018). Molecular Architecture of the Mouse Nervous System. Cell, 174(4), 999–1014.e22. 10.1016/j.cell.2018.06.021

