## Supplementary figures and images for "Glutamatergic and GABAergic synapses in the human spinal dorsal horn revealed with immunohistochemistry"

### Supplemental Video 1

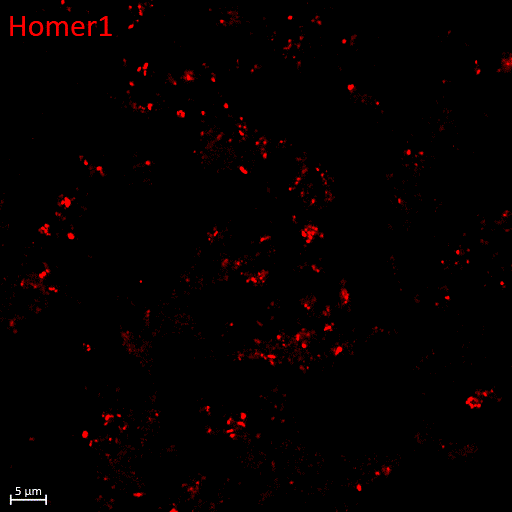

### Supplemental Video 2

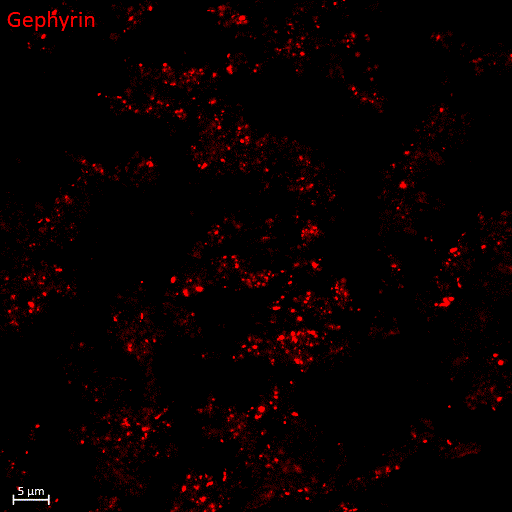
